# Specialized for the reach: Fruit picking and positional behavior favor a reach over a grasp phenotype for Geoffroy’s spider monkey (*Ateles geoffroyi* )

**DOI:** 10.1101/2025.05.15.654288

**Authors:** Ian Q. Whishaw, Jordan Dudley, Paulo Ramírez González, Evin Murrillo Chacon, Megan A. Mah, Fernando A. Campos, Filippo Aureli, Amanda D. Melin

## Abstract

The Geoffroy’s spider monkey (*Ateles geoffroyi*) has distinctive features, including a vestigial external thumb, long fingers and forelimbs, and a prehensile tail. To better understand how its derived morphology is used during natural foraging, we filmed wild spider monkeys well habituated to human observers in Sector Santa Rosa (SSR), Área de Conservación Guanacaste in northwestern Costa Rica. We analyzed frame-by-frame video recordings to examine the influence of this morphology on picking 14 fruit species. The spider monkeys’ most frequent reach strategy was a branch-withdraw (62%: 1,338 of 2,164 fruit items), in which they reached equally with either hand for a branch, hooked their fingers around it, and pulled it toward themselves to take the attached fruit by mouth. Less frequently they picked fruit by hand (466 observations) or only by mouth (360 observations). Hand and arm extension, with rotatory movements at the wrist, elbow, and shoulder, assisted reaching and transferring food to the mouth, and tail prehension further assisted extending reaches horizontally and ventrally into the tree canopy. To pick fruit by hand, they used tactile guidance and finger pad-to-palm grasps, mainly with the second digit (index finger), and rotatory arm and head movements to deliver fruit from the hand to the mouth. Fruit was grasped by the incisors and chewed with molars to release the endocarp for swallowing, with the exocarp sometimes ejected by spitting. The combination of branch-withdrawal, mouth grasping and postural extension via a prehensile tail constitutes a “reach” phenotype that contrasts with the “grasp” phenotype of the sympatric capuchin monkey. The behavioral and morphological commitment to a reach phenotype in spider monkeys supports the idea that the reach and the grasp have separate evolutionary histories and, with respect to spider and capuchin monkeys, contribute to niche partitioning.

“*Spider monkeys cast a distinct morphological silhouette – long scrawny arms and a snaky prehensile tail arching from a narrow pot-belly torso, topped by a small round head and blunt face. The commitment of this relatively large-bodied platyrrhine to a large-tree, upper canopy milieu and to ripe fruit foraging is seen throughout its skeletal and craniodental morphology.*” Rosenbert et al (2008).

## Introduction

The visual control of reaching has been of longstanding interest to evolutionary biologists and psychologists, who are interested in the evolution of skilled hand use in humans and its cognitive implications. Skilled forelimb use, such as that employed in eating by humans, has been suggested to be composed of a number of features, including the reach, the grasp, withdrawal to the mouth, posture, and laterality (Grant and Conway, 2019; Jeannerod, 1981; Jeannerod et al., 1995, 1998; Knecht, 2000; MacNeilage, 1987; Marzke, 1971. Nashner et al., 1985; Sartori et al., 2015; Whishaw et al., 2019, 2025). The modular structure of reaching has stimulated the suggestion that its components may have separate evolutionary histories (Karl and Whishaw, 2013; Whishaw and Karl, 2019). Comparative studies of the visual control of reaching in nonhuman primates provides evidence for phylogenetically related variation through which potential histories can be traced. Strepsirrhines, an early-branching suborder of primates, display visually guided reaching but not hand shaping when grasping or holding food (Peckre et al., 2019, 2023). By contrast, catarrhine primates, such as macaques (*Macaca*), feature a visually guided reach and grasp and visually mediated food handling (Hirsche et al., 2022; Macfarlane and Graziano, 2009; Marzke et al., 2015; Pouydebat et al., 2008; Scott, 2019) Among platyrrhine primates, capuchins excel in a visually guided grasp but do not display thumb/finger opposition, as used by catarrhine primates when grasping (Christel, 1993; Christel and Fragaszy, 2000). There is an extensive literature documenting the search for laterality in primates, and the study of spider monkeys has produced mixed findings (Boulinguez-Ambroise, 2022; Caspar, 2022; Hook-Costigan et al., 1996; Motes Rodrigo et al., 2018; Nelson et al., 2015a, 2015b), although there has been no study of spider monkey handedness in foraging. The present study was undertaken to add to the comparative understanding of feeding by studying the foraging of spider monkeys in the upper tree canopy.

The spider monkey, genus *Ateles*, is a large platyrrhine primate found across Mexico, Central, and South America (Mittermeier and van Roosmalen, 1981; van Roosmalen, 1985; Youlatos, 2002, 2008). *Ateles* are considered ripe fruit specialists, and the diet of *Ateles geoffroyi* consists of approximately 80% fruit from as many as 364 plant species (González Zamora et al., 2009; Scherbaum and Estrada, 2013). They prefer the softest ripe fruit, the seeds of which are defecated, with the resultant dispersal assisting the generation of new trees (Link and Di Fiore, 2006; Stevenson et al., 2002; Whitworth et al., 2019). They occasionally eat leaves, flowers, and slower-moving insects (Link, 2003). Their distinctive morphological features—including physique, limb joint hypermobility, prehensile lips, large spatulated incisors, and non-serrated molar teeth—have invited several explanations (Cant, 1986; Jenkins et al., 1978; Jungers and Stern, 1981; Mittermeier and van Roosmalen, 1981; van Roosmalen, 1985; Riba-Hernández et al., 2003; Youlatos, 2002, 2008). For example, the locomotion theory (Cant, 1986; Strier, 1992; Rosenberger, 1992; Youlatos, 2008) posits that brachiation with tail-assisted support aids movement between and within a patchily distributed ripe fruit tree habitat, while the diet theory (Rosenberger et al., 2008) proposes that obtaining ripe fruit located in a fine-branch, upper canopy milieu is central to their distinctive morphology.

Spider monkeys also feature a vestigial or absent external thumb (Melin et al., 2022; Saint-Hilaire, 1806; Tague, 1997), a feature that has not received the attention directed toward the capuchin monkey’s catarrhine-like thumb in studies of hand skills (Christel & Fragaszy, 2000; Costello & Fragaszy, 1988; Nelson, 2024; Melin et al., 2022; Spinozzi et al., 2004; Truppa et al., 2019, 2021; Whishaw et al., 2024a, 2024b). A thumb is essential for precision grasps involving opposition between the thumb and the fingers (Napier, 1956, 1962)—grasps proposed to be enabled by corticospinal projections onto motor neurons, which are rich in capuchin monkeys and catarrhine primates (Bortoff and Strick, 1993; Strick et al., 2021). In the limited study of the spider monkey neural systems (Chang and Ruch, 1947; Chico-Ponce de León et al., 2009; Lassek, 1943; Pubols and Pubols, 1972), the control of the hands has not been addressed. However, attention has been given to their prehensile tail, and the derived large brain that they share with other frugivores (DeCasien et al., 2017; Milton, 1993). Understanding spider monkey hand use for foraging may shed light on the structure of their visual control of reaching as well as the juxtaposition of their absent thumb and large brain. Possible outcomes might include enhanced fruit picking via tail-assisted reaching, impoverished fruit picking due to an absent thumb and impaired grasp, or one or more species-specific behavioral modifications that enable specialized fruit-picking skills.

## Methods

### Ethics Statement

This research adhered to the laws of Costa Rica, the United States, and Canada and complied with protocols approved by the Área de Conservación Guanacaste (R-SINAC-ACG-PI-027-18) (R-025-2014-OT-CONAGEBIO), by the Canada Research Council for Animal Care through the University of Calgary’s Life and Environmental Care Committee (AC19-0167).

### Study Population

The feeding behavior of *Ateles geoffroyi* was filmed in the Sector Santa Rosa (SSR), Área de Conservación Guanacaste (ACG) in northwestern Costa Rica (10.836°, -85.615°). The habitat is a seasonal tropical dry forest, where long-term study of spider monkeys has been ongoing, described in detail elsewhere (Asensio et al 2012; Melin et al 2020). The age and sex distribution of 34 Geoffroyi’s spider monkeys at filming were adult female (n=13), adult male (n=2), infant female (n=1), juvenile female (n=1), juvenile male (n=8), subadult female (n=6) and subadult male (=4). Videos was obtained from one animal as a subadult female and later as an adult female. Thirty of the spider monkeys were individually identified and 4 were not identified from the film clips. The animals appeared to be in good health and featured no disabilities that interfered with climbing, food grasping or other aspects of feeding. Filming consisted of short (7sec–10 min) continuous video samples following a published protocol with strict out-of-sight rules, such that recording was done when there was a relatively unobstructed view of the focal monkey’s feeding (Melin et al., 2022). Individuals were sampled opportunistically, based on visibility, but distribution of the observations among sex and age classes were generally representative of the population. Each film clip was labeled with the monkey’s ID for the purpose of identification.

### Video Recording

Video recording at 30 frames per sec (fps) provided *ad libitum* recordings of natural eating behavior and were collected using Lumix DC-G9, Sony FDR-AX53, and Olympus OM-D E-M1 camcorders.

### Food items

Spider monkeys were filmed eating fruits of fourteen plant species: *Doliocarpus dentatus* (ripe fruit diameter: 4 cm), *Bursera simaruba* (1 cm), *Coccoloba guanacastensis* (1 cm), *Dipterodendrum costaricensis* (2 cm), *Fictus ovalis* (1.5 cm), *Sciadodendron excelsum* (1.3 cm), *Karwinskia calderonii* (1.3 cm), *Guettarda macrosperma* (2.5 cm), *Ficus unknown* (3 cm), *Simarouba glauca* (1 cm), *Ficus cotinifolia* (1 cm), *Guazuma ulmifolia* (3 cm), *Spondias mombin* (4 cm), and *Bromelia plumieri* (6 cm).

### Video Analysis

We analyzed 134 video clips, which together comprised 4.48 hr of video. Each video clip provided many examples of hand use in stepping through the branches of the trees, reaching for food items or branches containing food items. Video recordings of spider monkeys were examined frame-by-frame using Quicktime 7.7 and Adobe Premier Pro software on an Apple computer by two observers (IQW and JD). Videos were scored using previously described methods (Hirsche et al., 2022; Peckre et al., 2023; Whishaw et al, 2024) and yielded an inter-scorer reliability coefficient of 0.96, based upon randomly selected videos representing approximately 10% of video cuts (Hallgren, 2012). Repeated scoring of the videos described feeding sequences in which a spider monkey reached for, used a hand or mouth to grasp, or withdrew a food item in the mouth. Because animals were reaching through leaves and adjusting posture, some component movements of some reaches were visible at times and others were obscured. The observable components were always scored.

### Behavioral Analysis

#### Positional Behavior

The scoring of spider monkey positional behavior is complicated by the many possible configurations of the limbs and tail (Cant, 1986). Here body position was scored on an 8-point scale (following Laird et al., 2022; Peckre et al., 2023) using an Eshkol-Wachman numeric-derived system (Golani, 1994; Whishaw et al. 2024). A score of “0” defined the horizontal long axis of an animal’s body orientation. Deviations of 45° were scored in steps in a clockwise order from 0 through 7; e.g., upright position = “6” and hanging straight down = “2”. Three aspects of behavior were scored:

1. *Grasping*. An animal received a positional score at the point that it grasped a food item whether by hand or mouth.
2. *Consumption*. An animal received a postural score at the point that the food item was taken by the mouth directly or from the hand.
3. *Tail support*. Tail support was scored as “1” if it was judged that the monkey would fall if tail support was lost. If positional behavior did not depend upon tail support a score of “0” was given.

#### Food-purchase movement

Four types of reaching and laterality were documented by counts:

*Grasp-withdraw*. A grasp-withdraw reach involved a single hand advancing to grasp a food item from a branch to bring it to the mouth.

*Branch-withdraw*. A branch-withdraw reach involved grasping a branch that was then brought toward the mouth so that a fruit item could be taken from the branch with the mouth.

*Mouth-withdraw*. A mouth-withdraw reach involved the mouth taking fruit items without the involvement of the hands.

*Inhand-withdraw*. An inhand-withdraw involved a food item held in the hand being brought to the mouth.

*Hand laterality.* Use of the left or the right hand for each of the above four reach/withdraw types were counted to determine individual and population hand asymmetry (Hook-Costigan and Rogers, 1996; Motes Rodrigo et al., 2018; Nelson et al., 2015).

#### Mouth handling

The way in which the mouth grasped food items from the hand was documented with counts of occurrences (Hirsche et al., 2022):

*Incisor grasp*. An incisor grasp involved the mouth opening and grasping a food item with the incisor teeth.

*Premolar grasp*. A premolar grasp involved the mouth opening and grasping a food item with the premolar teeth.

*Chewing*. Repeated mouth movements occurring following taking a food item by mouth were scored as chewing.

*Spitting*. If upon chewing a fruit item the monkey expelled some part of the fruit from its mouth, the act was scored as spitting.

*Fruit sucking*. If a monkey grasped a fruit item by mouth and chewed it without picking it from the stem, the act was scored as fruit sucking.

#### Component movements of reaching

*Head movement*. The head contribution to a withdraw-to-eat in reaching for a food item was scored on a 5-point scale (Hirsche et al., 2022; Peckre et al., 2023):

*Score 0*. The head was advanced toward the food item and the food item was grasped with the mouth.

*Score 1*. The nose was placed near the food item, but the item was grasped by hand and brought to the mouth by hand.

*Score 2*. The hand holding the food item and the mouth were brought toward each other such that the withdraw-to-eat movement was accomplished equally by the hand and mouth.

*Score 3*. Most of the withdraw-to-eat was accomplished with the hand holding the food item with only a small orienting movement made by the mouth toward the hand.

*Score 4*. The head was not advanced toward the food item, or even withdrew, as the hand brought the fruit toward the mouth.

*Hand movements.* The following hand movements were scored:

*Pronation.* A hand movement that involved hand rotation such that the palm faced downward was designated as pronation.

*Supination.* A hand movement that involved hand rotation such that the palm faced upward was designated a supination.

#### Statistical Analyses

Counts of behaviors including posture, grasping type, engage and disengage, blinking and eye direction are reported as the percent of the number of observations that were made. The relationship between the duration of the immediate withdraw-to-eat and in-hand withdraw-to-eat movements to head-engage, eye engage, and disengage duration were assessed using the Pearson product-moment correlation and compared with Chi-square tests. Group comparisons were made using a generalized linear model repeated measure analysis of variance on SPSS (29.0.1.1). Laterality measures were made using t-tests with the comparison variable being chance occurrence.

## Results

### General foraging observations

Video footage demonstrated spider monkeys are highly adept in moving through the trees, including the distal branches of trees (Figure 1). Feeding principally occurred in the canopy, in which the monkeys harvested fruit from the terminal canopy branches. The spider monkeys displayed a rich array of positional behavior and food purchase strategies to access food on every location on a tree. Given that the data were obtained from the spontaneous behavior of the animals in trees, the movements of reaching were associated with the movement of the animals themselves, the movements of the branches produced by the animal’s positional changes, the wind, and movements of other animals. Behavioral observations were scored despite the complexity of substrate and positional movements. There were some occurrences of reaching acts associated with bringing a food item to the nose for sniffing, grasping a food item and not picking it, and grasping fruit that was then dropped (Milan et al., 20129), but these were not counted.

**Figure 1.**
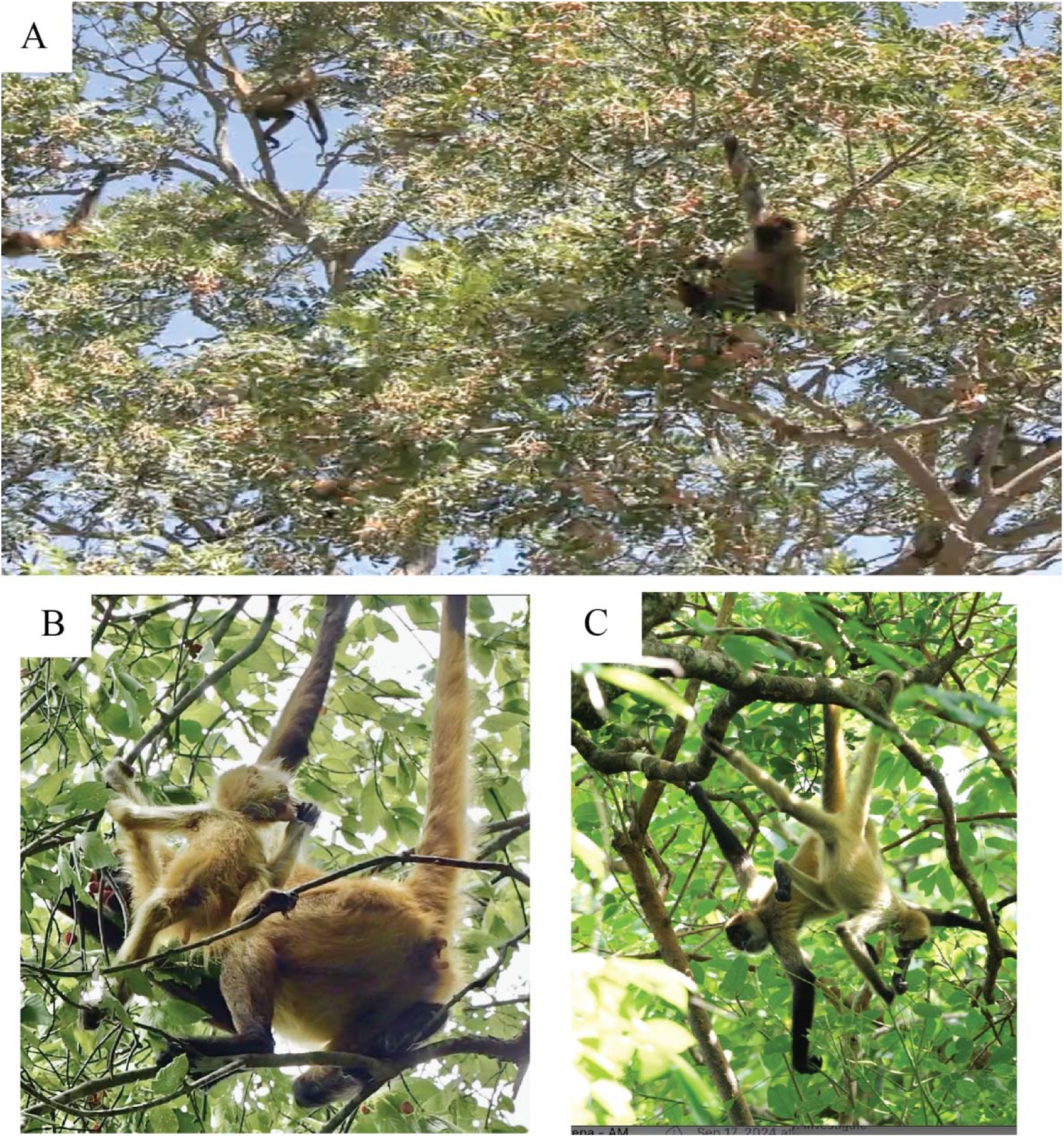
**Spider monkeys (Ateles geoffroyi) foraging**. *Note*. A. Hungria, an adult female, foraging on the distal branches of a *Simarouba glauca* fruit tree. B. Inverness, an adult female with an infant, foraging in the branches of a *Guettarda macrosperma* tree. C. Inverness (background) with offspring, now a subadult (foreground) foraging on the leaves of a *Guettarda macrosperma* tree.

#### Positional behavior and prehensile tail use

Figure 2 (Video 1) summarizes positional behavior and the corresponding use of the tail for support. Figure 2A illustrates the scheme for the numerical scoring (0-7) for position based on the long axis of the body in the vertical plane.

**Figure 2.**
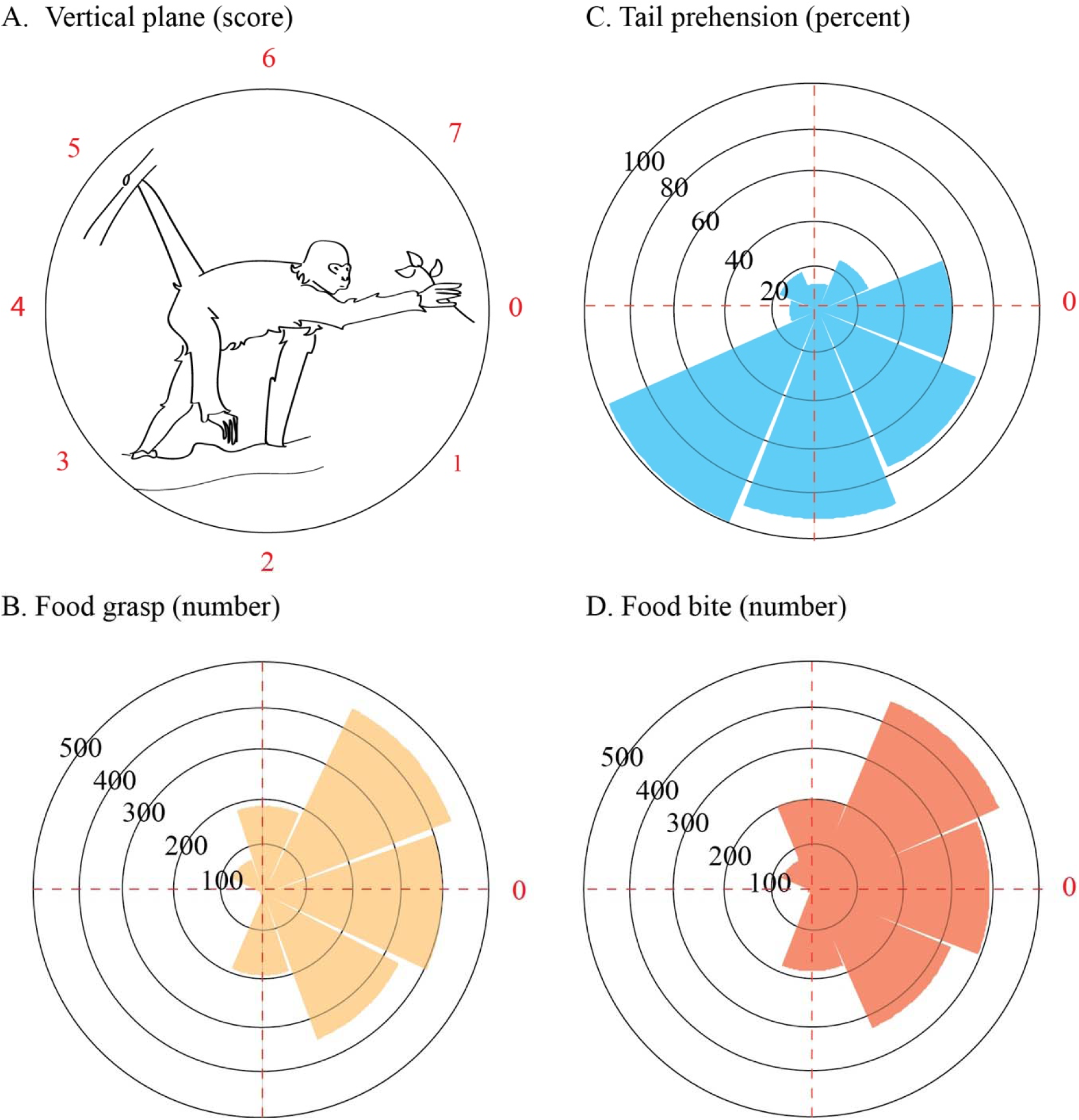
**Positional behavior and tail use for grasping and eating fruit by spider monkeys***. Note*. A. Scoring system (from 0-7) for positional behavior that scores the long axis of the body in deviations of 45°. B. Percent of body positions used by spider monkeys that depended on tail support during reaching for a food item. C. The number of times spider monkeys adopted a body position as they reached for a fruit item. D. The number of times a spider monkey adopted a body position as they placed a fruit item in the mouth. The correlation between the body position used while reaching for fruit and the body position used while subsequently placing fruit in the mouth was r=.940, indicating that posture was not changed between grasping (C) and placing fruit in the mouth (D).

**Figure 3.**
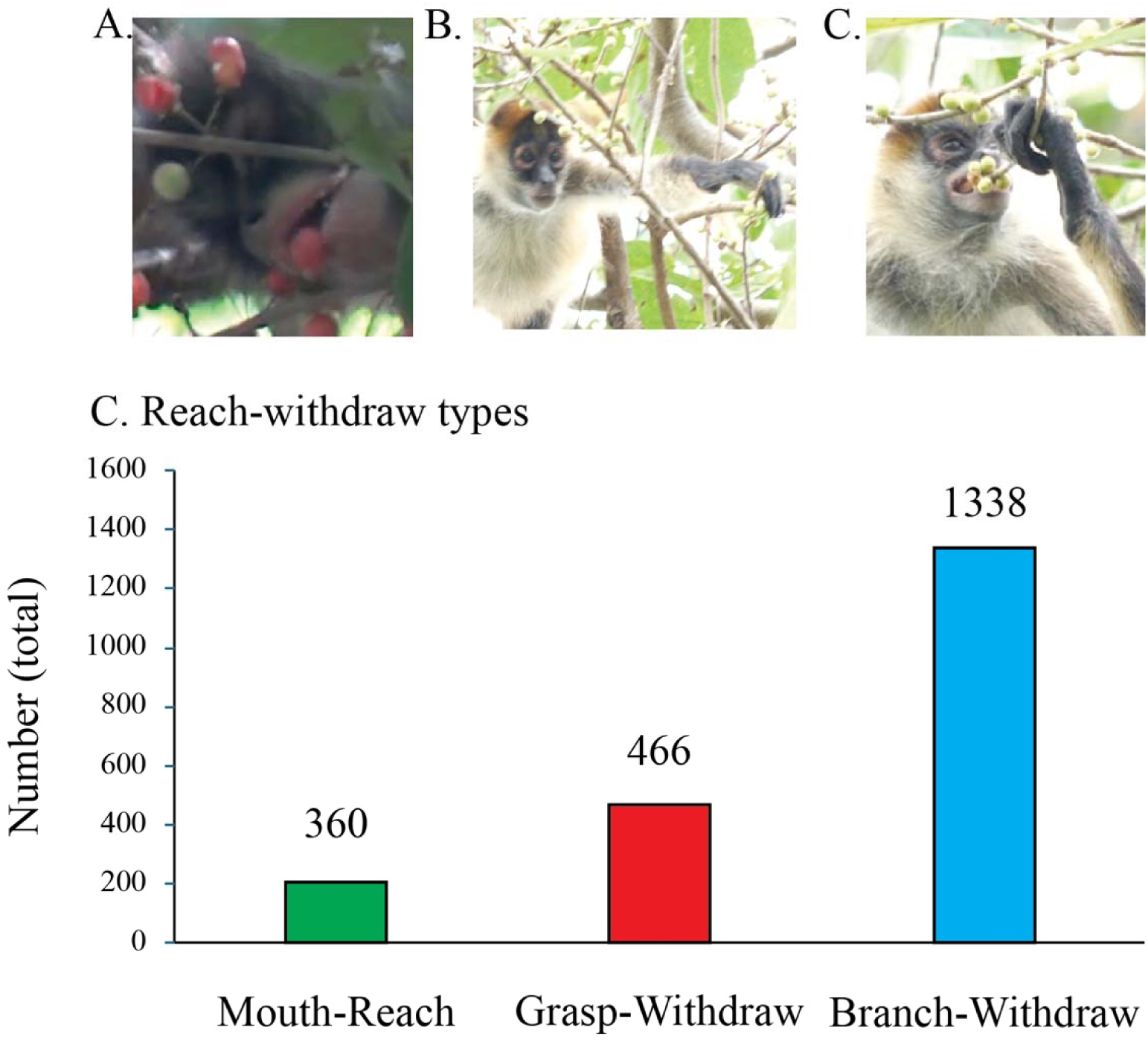
**Three fruit purchase strategies used by Ateles Geffroyii for obtaining fruit**. *Note*. A. Mouth-withdraw: a spider monkey named Kenya displays a mouth-withdraw in which an individual fruit of *Doliocarpus dentatus* is taken by mouth. B. Hand-withdraw: a spider monkey named UN displays a hand-withdraw in which a single *Fictus cotinifolia* fruit is taken with the hand. C. Branch-withdraw: the spider monkey UN displays a branch-withdraw in which a branch containing *Fictus cotinifolia* is pulled by hand toward the mouth so that the fruit can be picked by the mouth. D. The frequencies of three types of fruit purchase out of a total of 2,164 retrieval events observed, illustrating that branch-withdraws were about twice as frequent as grasp-withdraw reaches and mouth-reaches combined.

Figure 2B illustrates the percentage of positions in which a spider monkey depended upon tail support when grasping a food item. In all, 770 of 1597 reaches (48 %) were dependent on tail support. The use tail support in relation to position was similar across all animals, as indicated by a significant group position effect, F(7,210)=2.36, p=.024, with no significant difference in tail use in adults vs juveniles/subadults, F(1,30)=0.78, .782, or males vs females, F(1,30)=.632, p=.433.

Figure 2C illustrates the distribution of the number of positions adopted by the spider monkeys as they grasped 1,597 fruit items. There was a significant group effect of position, F(7,210)=11.2, p<.001, in which sit, horizontal, and slightly inclined were the most frequently used positions. There was no significant difference in the distributions of positions adopted by adults vs subadults/juveniles, F(1,30)=1.02, p=.319, or males vs females, F(1,30)= .26, p= .616. Correlation of the distribution between positions used for the grasp and positions when placing the fruit in the mouth was highly significant (was r=.940), showing that there was little change in an animals position between picking a fruit item and then placed in the mouth (Figure 2C fs 2D).

#### Mouth-withdraw

Mouth-withdraw movements (Video 2) were used for taking all species of fruits. There was no correlation between fruit size and the incidence of mouth-withdraw movements (Pearson correlation: r =.320, F(1,14)=1.3, p=.270). Nevertheless, some fruit species were preferentially taken directly by mouth. The highest incidence was associated with picking the fruit *Sciadodendron excelsum,* which grows in upward facing clusters on relatively large branches. The only strategy observed for taking this fruit was with the mouth, usually from a position with the head located above the fruit cluster (18.5% of all mouth withdraw observations). The second highest incidence (15.6% of all mouth-withdraw observations) of the use of mouth-withdraw was with the fruit *Doliocarpus dentatus*. On instances where the monkeys did pick this fruit by mouth, they could be observed to spit out the shell after chewing, as is shown in the first sequence in Video 2. There were also instances in which they bit and sucked *Doliocarpus dentatus* to obtain the endocarp while leaving the endocarp hanging on the vine. A representative sequence of fruit bite/suck is shown in the last two sequences of Video 2.

#### Grasp-withdraw

Grasp-withdraw consists of reaching with a hand to pick fruit and Figure 4 (Video 3) illustrates a representative grasp-withdraw movement. The movement consisted of: (1) directing gaze to the food item, (2) advancing a hand (Figure 4A), (3) grasping the fruit (Figure 4B), (4) withdrawing the fruit to the mouth and (5) biting the food with the incisors (Figure 4C). Figure 4D gives a kinematic representation of a grasp-withdraw associated with obtaining a small food item, *Ficus cotinifolia*. The figure illustrates that the monkey directed gaze to the target at about the time that a reach was initiated, then the monkey grasped the food item after vision was disengaged, and finally the monkey withdrew the fruit to the mouth to grasp it in the incisors.

**Figure 4.**
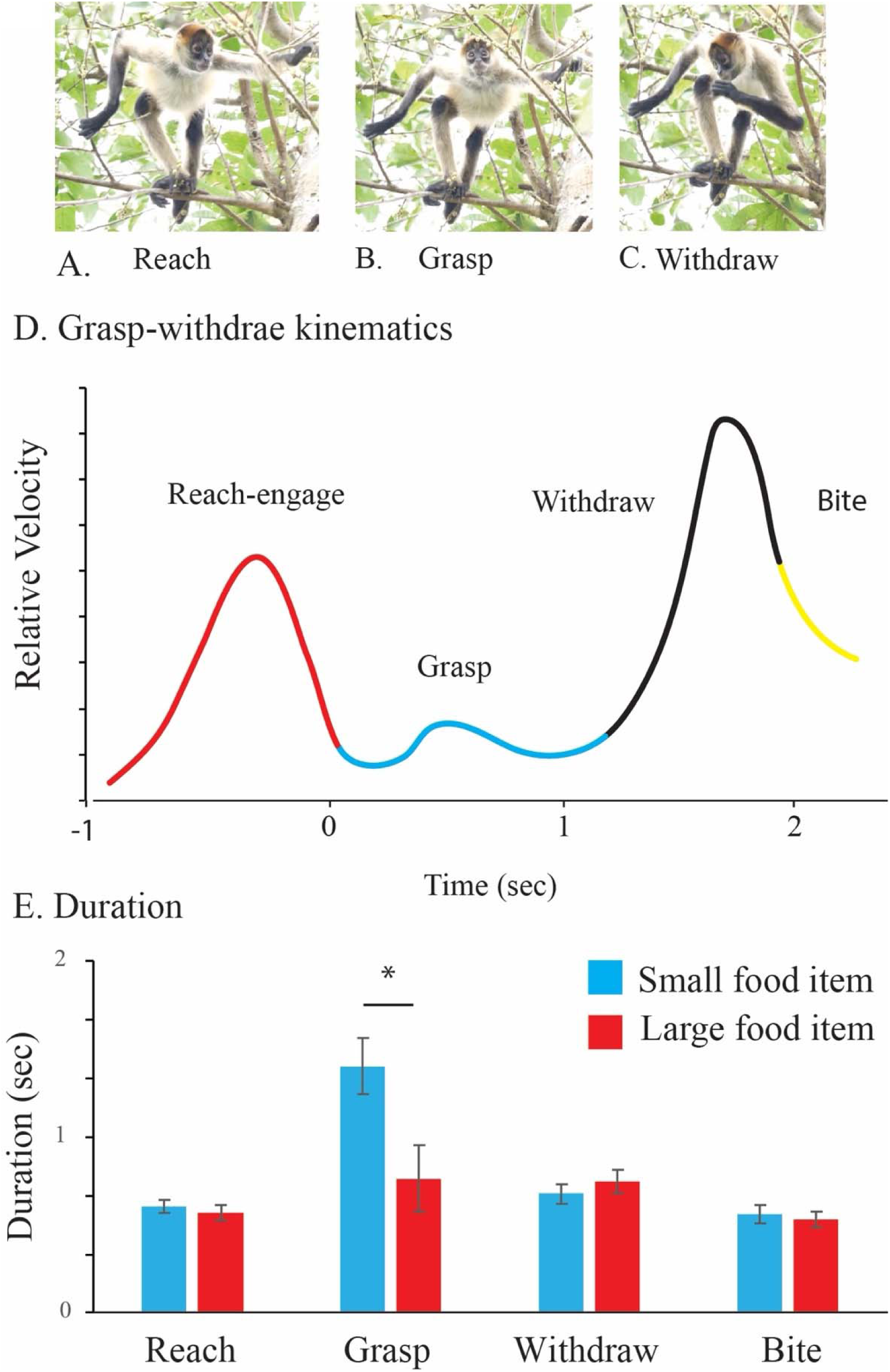
**Grasp-withdraw reach made by a spider monkey named UN eating the fruit Ficus cotinifolia**. *Note*. The grasp-withdraw movement involves: A. a reach with a hand to take a fruit item, B. picking the item, C. withdrawing the item to the mouth. D. Kinematic representation of a grasp-withdraw for a *Ficus cotinifolia* fruit illustrating the duration of the reach, the grasp, and the withdraw to the mouth. E. Durations (mean±se) of grasp-withdraw movements for small fruit items (<1.5 cm) vs large fruit items (>1.5 cm). The grasp duration associated with small fruit items was longer than for the large fruit items, *=p<.05.

An analysis of the relative duration of each grasp-withdraw component was made from 108 grasp-withdraw movements in which all components could be observed in the video sequence. Figure 4E illustrates the comparison of the duration of the reach, grasp, withdraw and bite movements for the smaller fruit items (>1.5 cm, n=45) vs the larger fruit items (<1.5 cm, n=63). An ANOVA indicated that the time taken to purchase the smaller fruit items was significantly longer than the time taken to purchase the larger fruit items, F(1, 106) = 4.07, p=.046, *^2^* = .04. There was a significant interaction between duration for the different movement components in relation to fruit size, F(1,106)=5.93, p<.001, *^2^* = .04. Follow-up Newman-Kules tests indicated that the time taken to make the grasp of smaller fruit items was longer than the time taken to grasp larger fruit items (p=.020). For the other movement components, the reach, withdraw, and bite, the durations were not different in relation to fruit size. Observation of how grasps were made suggested that the longer times taken to make a grasp of a small fruit item was due to the time taken to position the hand in relation to the fruit so that the fingers could close on it (see below).

#### Branch-withdraw

Figure 5 (Video 4) illustrates the components of a branch-withdraw when obtaining *Ficus cotinifolia,* a small fruit item. Components are grasping a branch containing a target fruit with one hand (Figure 5A), withdrawing the branch toward the mouth (Figure 5B), then reaching with the mouth to take a target fruit item from the branch with the incisors (Figure 5C). A kinematic representation of this sequence is shown in Figure 5D, in which the knuckle of the second digit of the hand holding the branch and the middle of the upper lip are digitized. The kinematic representation shows that as the branch is withdrawn, the head orients toward the fruit, the fruit is visually engaged, the mouth is advanced to take the fruit, while at the same time disengaging vision, and then using the incisors take the fruit.

**Figure 5.**
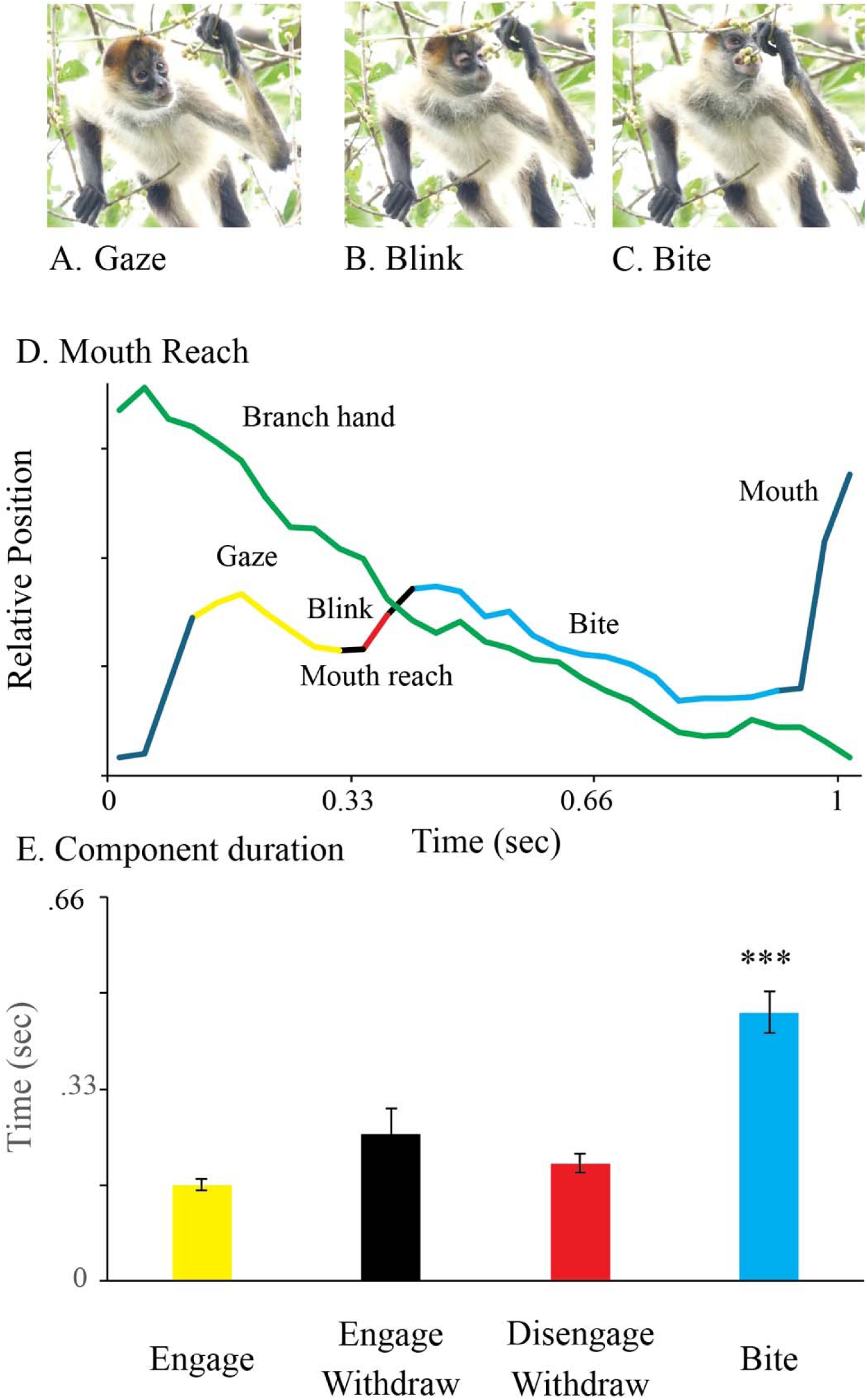
**Branch-withdraw movement made by a spider monkey UN eating the fruit Ficus cotinifolia**. *Note*. The branch-withdraw movement involves: A. Directing gaze toward the fruit as the mouth begins its advance to take the fruit; B. Blinking with gaze disengagement as the mouth is moved toward the fruit; C. Raising the head to bite the fruit with the incisors to pick it from the branch. D. Kinematic representation of a branch-withdraw for a *Ficus cotinifolia* fruit illustrating the movement of the hand grasping the branch (the knuckle of the second digit is digitized) and the movement of the mouth (middle of the upper lip is digitized) to grasp the fruit. For the bite, the movement of the fruit and the mouth are roughly parallel. E. Duration (mean±se) of the components of branch-withdraw movements of the mouth to take fruit obtained from 28 branch-withdraw reaches. The component that takes the longest to complete is the bite, ***=P<.001.

It was difficult to observe all gaze features associated with branch-withdraw movements because when the face was inclined, the eyes were obscured against the dark facial fur of the spider monkeys. Nevertheless, 27 branch-withdraw movements were identified in which the eyes were clearly visible throughout the movement sequence. On 23 of these occasions, the monkeys disengaged gaze with a blink before opening the mouth to grasp the fruit with the incisors. Figure 5E illustrates the average duration of branch-withdraw component movements. An ANOVA on component durations of gaze, withdraw with gaze, withdraw without gaze, and bite with the incisors gave a significant effect, F(1,26)=49.9, p<.001, *_η_*2=.66. Newman-Kules tests indicate that the bite duration (time from the lip touching the food to its removal from the hand) was the component that took the longest to complete, p<.001.

Figure 6 illustrates the relative use of branch-withdraw movements in relation to the total number of fruit grasp movements for the 14 different fruits. Figure 6A illustrates the percent of branch-withdraw movements for each fruit. Figure 6B shows there was a significant correlation between fruit size and the incidence of a branch-withdraw, r=.69, F(1,12)=11.4, p=.006, with branch-withdraw more likely to be used for small fruit items.

**Figure 6.**
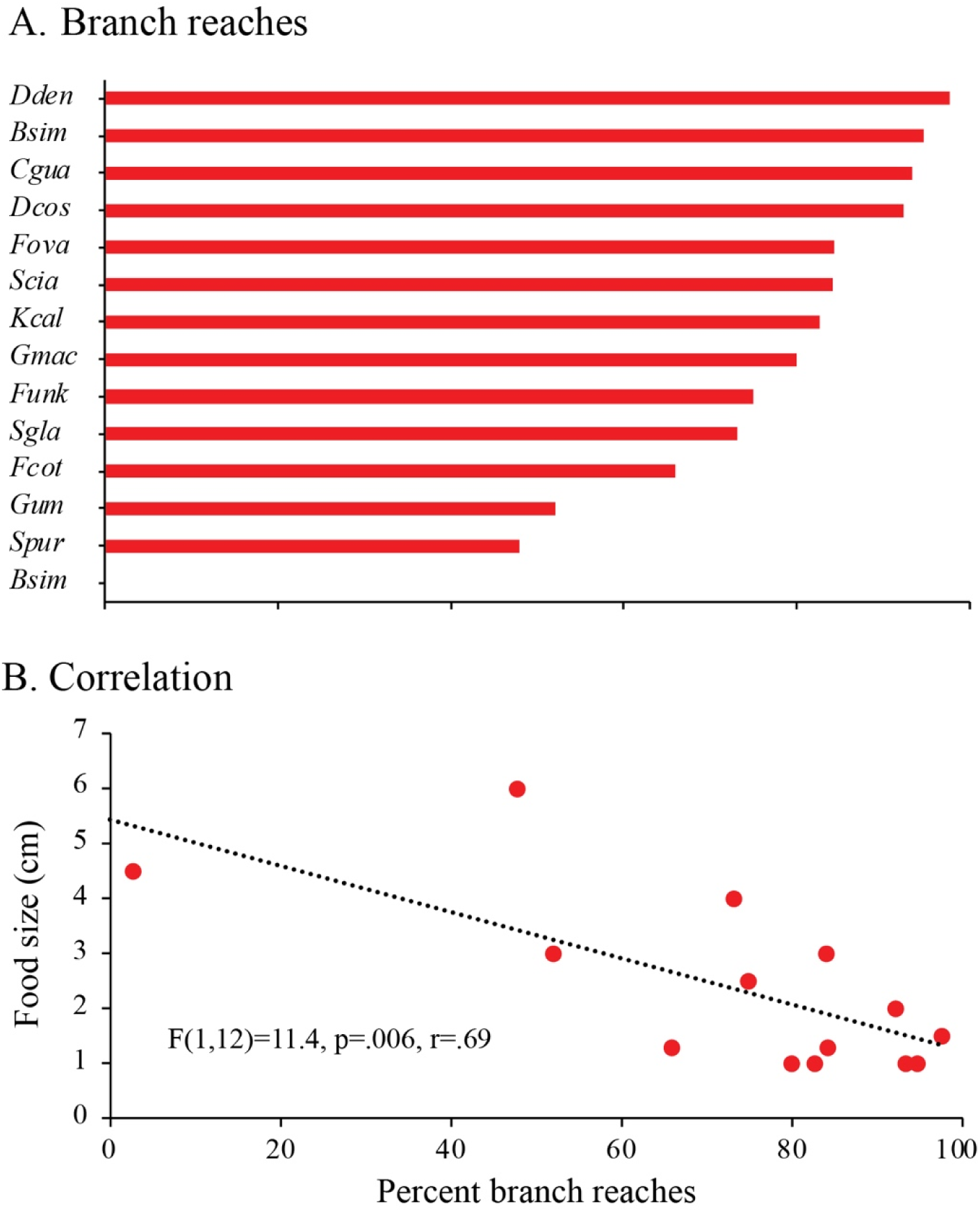
***A. The percent of all fruit retrieval events that were branch-reaches as a function of fruit species***. *Note.* A. The probability of using a branch-reach strategy of fruit retrieval as a function of fruit species diameter. B. Correlation between branch-reach strategy and fruit species size. The smallest fruit were the most likely to be associated with a branch-retrieval strategy. Food items were: *Doliocarpus dentatus* (*Dcec*), *Bursera simaruba* (*Bsim*), *Coccoloba guanacastensis* (*Cogu*), *Dipterodendrum costaricensis* (*Dcos*), *Fictus ovalis* (*Fova*), *Sciadodendron excelsum* (*Sexc*), *Karwinskia calderonii* (*Kcalcm*), *Guettarda macrosperma* (*Gmac*), *Ficus unknown* (*Fun*), *Simarouba glauca* (*Sgla*), *Ficus cotinifolia* (*Fcot*), *Guazuma ulmifolia* (*Gulm*), *Spondias mombinplumieri* (*Bplum*), *Bromelia plumieri* (*Bplum*).

On 32 of the 1338 branch-withdraw movements observed, the food was taken with the other hand and not the mouth. For these observations, however, it was difficult to determine the extent to which a monkey was withdrawing a branch as opposed to holding onto it for support.

In sum, these results suggest that for grasp-withdraw movements, the smallest fruit items are the most difficult to grasp and the most difficulty to transfer to the mouth.

#### Inhand-withdraw movements

Inhand-withdraw movements, in which a fruit held in the hand is brought to the mouth (Figure 7, Video 5), were only observed during eating of the fruit *Guazuma ulmifolia* (n = 155 representing an average of 21.1 observations in each of 7 spider monkeys. *Guazuma ulmifolia* is a 2-3cm long fruit featuring a hard exocarp that surrounds an endocarp of seeds, which are the targets of eating (Manríquez-Mendoza et al., 2011; Pereira et al, 2019).

**Figure 7.**
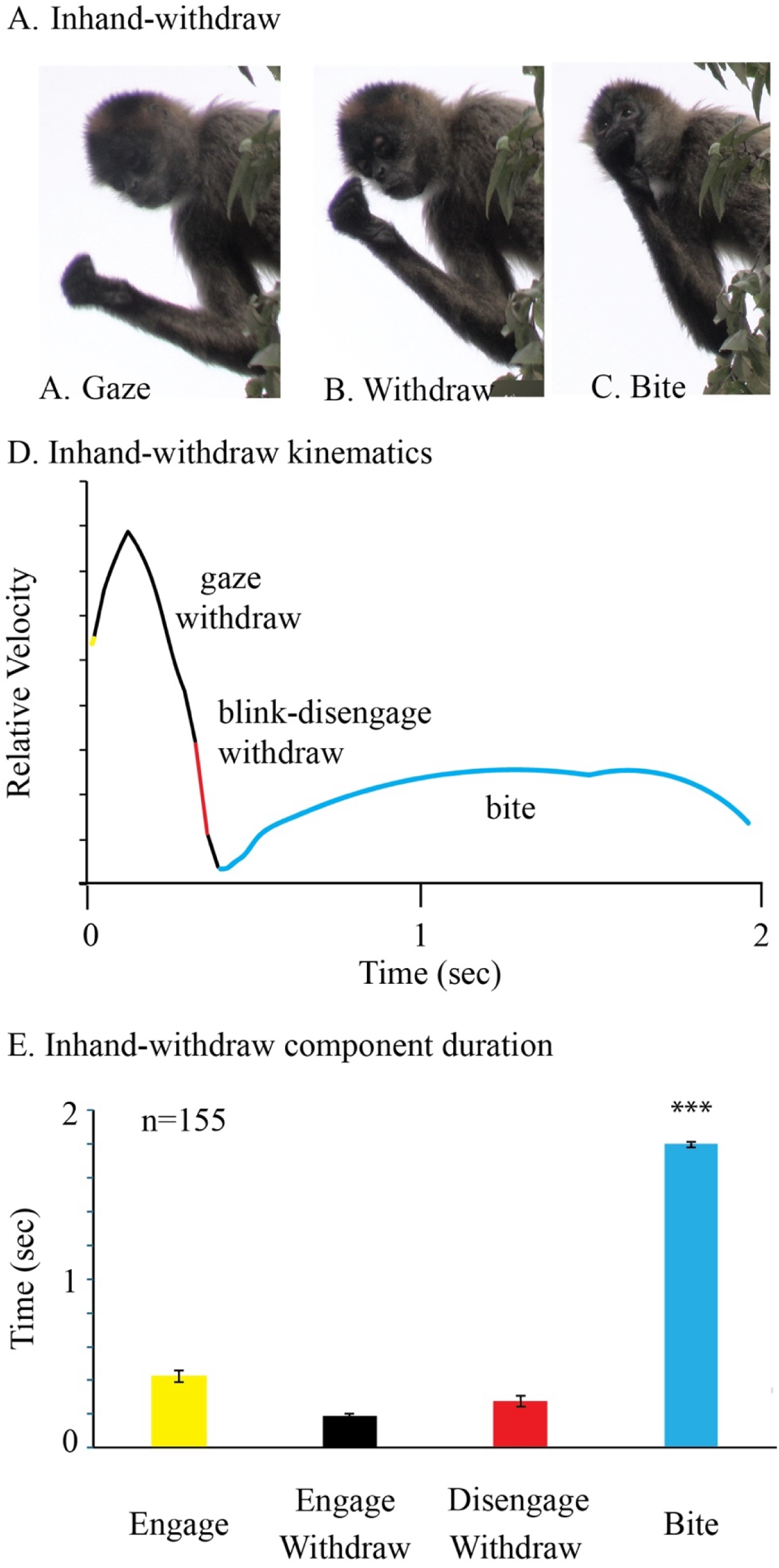
**Inhand-withdraw by a spider monkey named Ajka eating the fruit Guazuma ulmifolia Note**. The fruit is held in the hand and brought to the mouth for eating in each inhand-withdraw, which consists of: A. Directing gaze toward the fruit held in the hand; B. Withdrawing the fruit toward the mouth while maintaining gaze on the fruit; C. Disengaging gaze from the fruit and concurrently raising the head so that the mouth is in a horizontal orientation to receive the fruit from the hand. D. The relative velocity and duration of the hand’s component movements of reach grasp, bite during an inhand-withdraw that brings the hand to the mouth for chewing. Note: the subject blinks, as gaze is disengaged from the fruit, and the bite duration is relatively long with the hand moving the fruit between the incisor and premolar teeth for chewing. Bottom, E. Relative duration (mean±se) of the component movements of inhand-withdraw movements observed in 7 spider monkeys eating the fruit *Guazuma ulmifolia.* Note: the bite takes relatively longer than the other movements, ***=P<.001.

Figure 7 shows a representative inhand-withdraw movement, which consists of first looking at the fruit (Figure 7A), then bringing it to the mouth under gaze (Figure 7B), and then disengaging vision while completing the withdraw movement to the mouth for biting (Figure 7C). A representative kinematic description of an inhand-withdraw is shown in Figure 7D, illustrating its four components: (1) gaze directed at the fruit held in the hand, (2) gaze during the first portion of withdraw, (3) gaze disengage during the second part of the withdraw and (4) grasping with the incisors and chewing with the premolar teeth. Of the 155 inhand-withdraw movements, the eyes were clearly visible in 97 movements, and of these, 96 were associated with a blink at gaze disengage. Associated with the gaze disengage, the head was raised so that when the fruit reached the mouth, the mouth was in a horizontal orientation. After chewing the fruit with the premolars, the spider monkeys spat out the exocarp and then swallowed the endocarp.

Figure 7E illustrates the relative duration of each component movement, showing that the movement component with the longest duration was biting and chewing the fruit (F(1,153)=193.9, p<.001, _η_2=.560).

#### Population and individual handedness

Figure 8 summarizes comparisons of left-hand vs right-hand use for picking fruit by 30 spider monkeys. Analysis of the frequence of hand preference use for picking and handling fruit gave support for individual left or right hand preferences but little support for a population hand-preference. Analysis of left and right hand use, assessed with t-tests for correlated groups, gave no significant difference for grasp-withdraw movements, t(29)=.18, p=.850. Individual animals did show preferences in the use of one or the other hand as shown by scatter plots of handedness scores (Figure 8A). Analysis of left vs right hand use for reaching for branches as a part of branch-withdraw movements did produce a small left hand preference, t(29)=2.3, p=.030, but again there were large individual differences in hand preferences (Figure 8B). Comparisons of hand preference for inhand-withdraw from the 7 spider monkeys eating *Guazuma ulmifolia* gave no significant preference in use of either hand, t(6) =12., p=.260. When the incidence of left and right hand use was summed for the three types of hand movement, in relation to the 14 different fruit species, there was no left vs right preference, F(1,13)=1.9, p=.08, although there were individual preferences, as illustrated by the scatter plot (Figure 8C).

**Figure 8.**
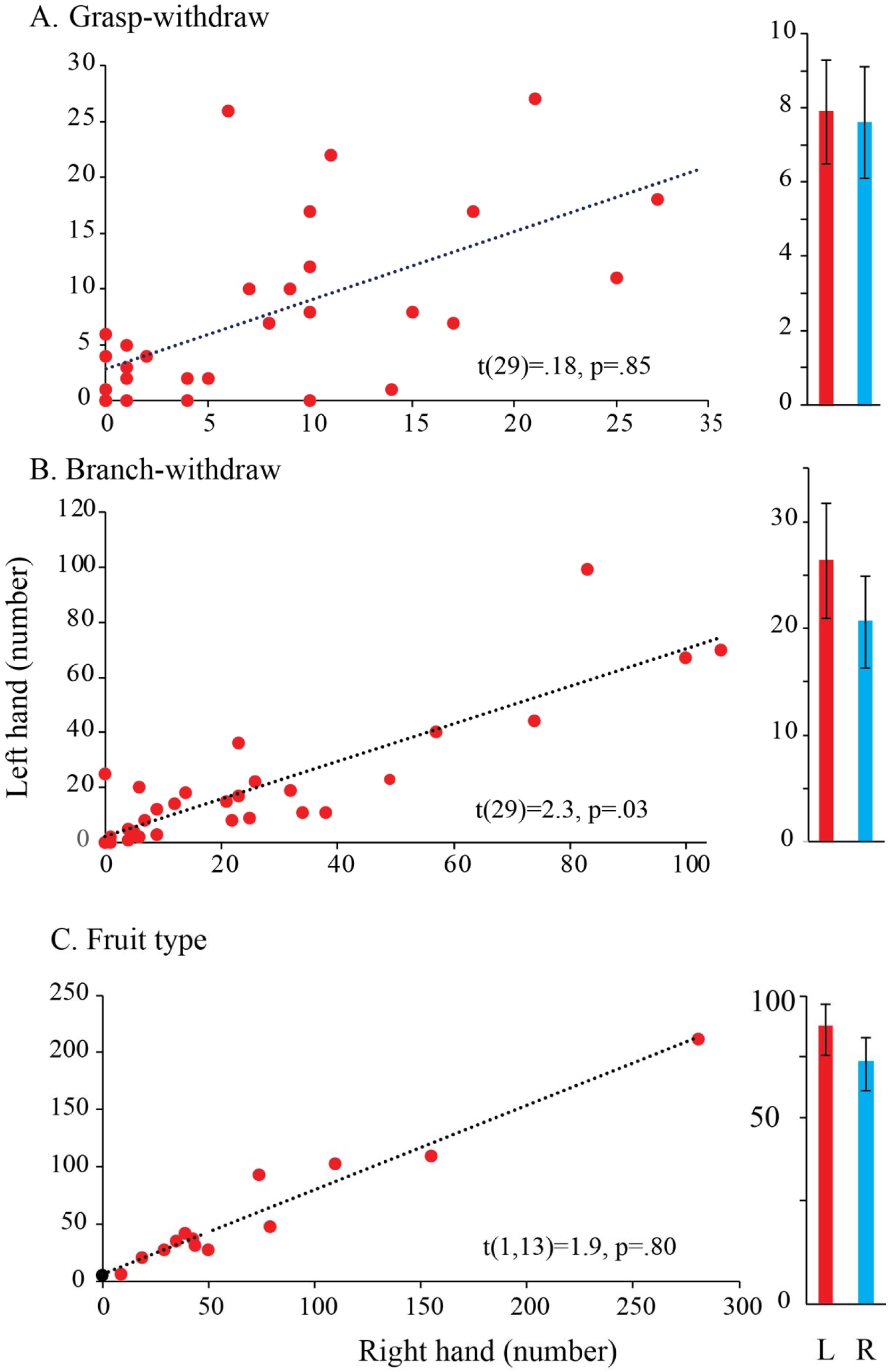
**Relative use of the left and right hands of individual Ateles geoffroyi**. *Note*. Scatter plots present left vs right hand use for each individual animal observed eating. Bar graphs give group left vs right hand use (mean±se). A. Grasp-withdraw reaches, in which a fruit item is taken by a hand. B. Branch-withdraw reaches, in which a branch is grasped in one hand and brought toward the mouth with the fruit taken from the branch by the mouth. C. Relative left vs right hand use for all reaches in relation to each of 12 different kinds of fruit species. The spider monkeys displayed very small to no preference for using either the left or right hands when purchasing fruit.

#### Components movements of food acquisition

Associated with the use of different food reaching strategies, the spider monkeys made different use of component movements. including use of the mouth for picking, the incisors grasping, the molars in processing fruit and rotatory movements of the hands and mouth for grasping and transferring fruit from the hands to the mouth. These are described in turn.

#### Head movement with reaching

Figure 9A illustrated two different head movements related grasping food with the incisors. The head was advanced to grasp fruit associated with branch-withdraw and it was retracting to accept food associated with grasp-withdraw. Figure 9B summarizes the 5-point score based on 565 branch-withdraw, 234 grasp-withdraw, and 219 inhand-withdraw movements. The ANOVA of head movement direction associated with withdraw types was significant, F(2,1012)=327, p<.001, η^2=^.86.. Follow-up Neuman-Keul tests (p<.001) confirmed that branch-withdraw movements were always associated with the head moving to take the fruit, whereas grasp-withdraw and inhand-withdraw movements were always associated with the head moving away from the fruit as the hand brought the fruit to the mouth.

**Figure 9.**
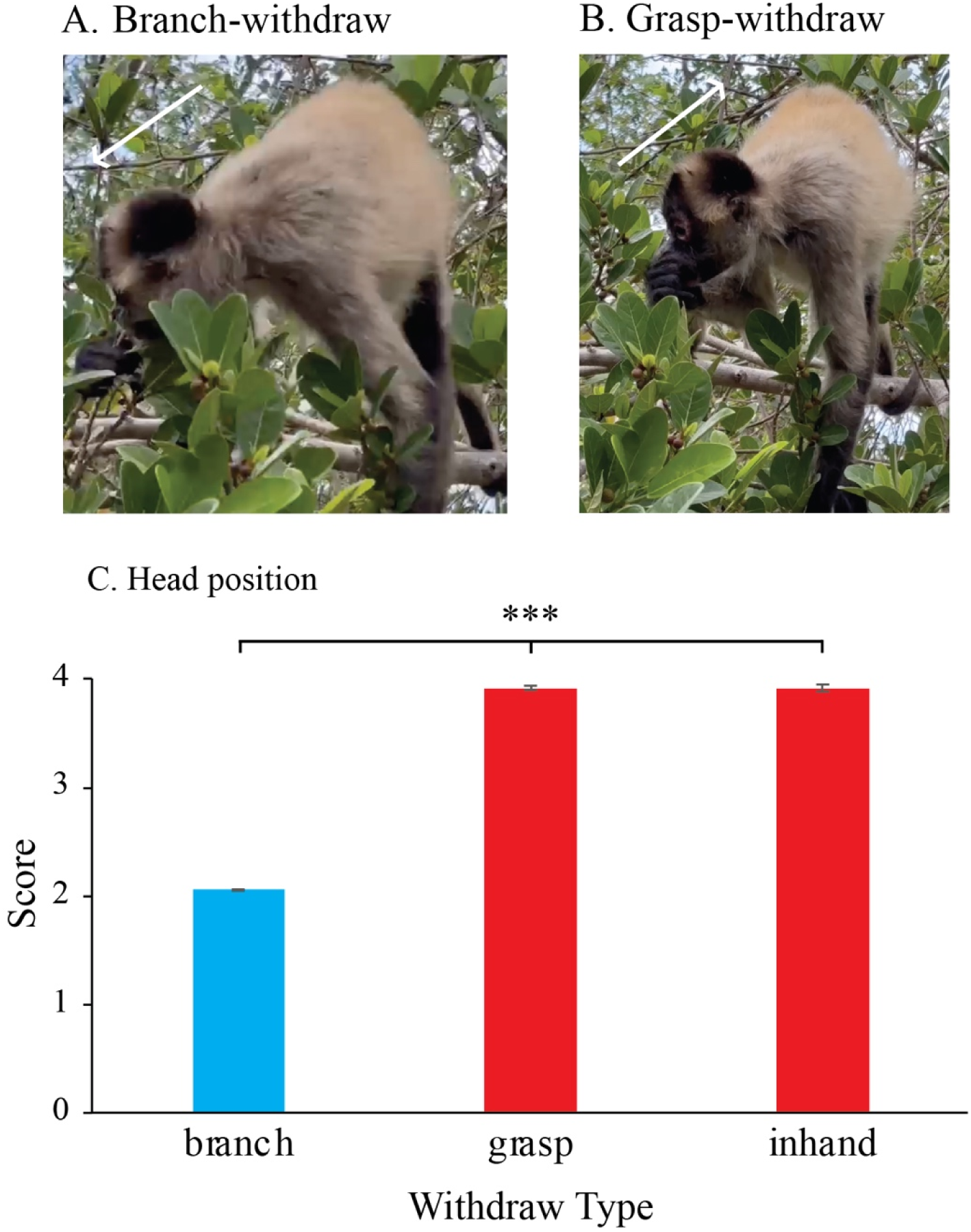
**A spider monkey called Quetzaltenango displays two kinds of head movements related to fruit purchase**. *Note.* A. Branch-withdraw reach for the fruit *Fictus cotinifolia* involves head advance to take the fruit by mouth. B. Grasp-withdraw movement for the fruit item involves raising the head, moving the snout away from the hand as it approaches the mouth. C. Scores (mean±se) of head movements for all spider monkeys displaying branch-withdraw, grasp-withdraw, and inhand-withdraw movements. Note: The head is advanced to take fruit into the mouth only when making branch-withdraw food purchases, ***=P<.001.

#### Fruit chewing

An example of fruit chewing is shown in Video 6. Of the 863 observations in which there was a clear view of a monkey placing food in the mouth, 838 featured the spider monkeys chewing the fruit. On 25 of the remaining food placements, in the mouth was obscured by head turning or leaves. Theus, the spider monkeys chewed fruit of all sizes as confirmed by a nonsignificant correlation of chewing with fruit size and type, r=.17; r (1,12) = 0.180, p=.54. For larger fruit items, such as *Spondias purpurea*, a plum-sized fruit the size of a small plum, a spider monkey were also seen to raised the head as chewing progressed, to seemingly aid in swallowing (Video 7). For some other fruit items, notably *Guazuma ulmifolia* and *Ficus cotinifolia*, both of which appeared to have fruit seeds enclosed in a tough exocarp, chewing appeared to be directed to removing the exocarp, which was then ejected by spitting. Occasionally after taking a fruit item into its mouth, the spider monkey simply spat the fruit out, suggesting that in addition to sniffing, taste was also used to assess edibility/ripeness (Melin et al, 2019). When harvesting *Doliocarpus dentatus*, spider monkeys were observed to pick the fruit by mouth and spit out the shell after chewing as well as to chew the fruit on the vine, leaving the shell still attached to the vine.

#### Hand supination and pronation

Figure 10 summarizes rotation of the hand associated with 792 branch-withdraw grasps and 562 grasp-withdraw grasps. When grasping a branch, the hand was pronated on 74% of branch-withdraws; that is, palm-up in relation to the target (Figure 10A). When grasping a fruit item, the hand was pronated on 63.6% of grasp-withdraws (Figure 10B). In addition, with respect of the direction of reaches, 66% of downward reaches featured pronation, 63% of horizontal reaches featured pronation, and 78% for upward reaches featured pronation. In all, although pronation was favored as a reach orientation, the choice of hand orientation seemed opportunistic, that is; influenced by the location of the target in relation to the monkey, and not to a constraint on hand rotation for grasping.

**Figure 10.**
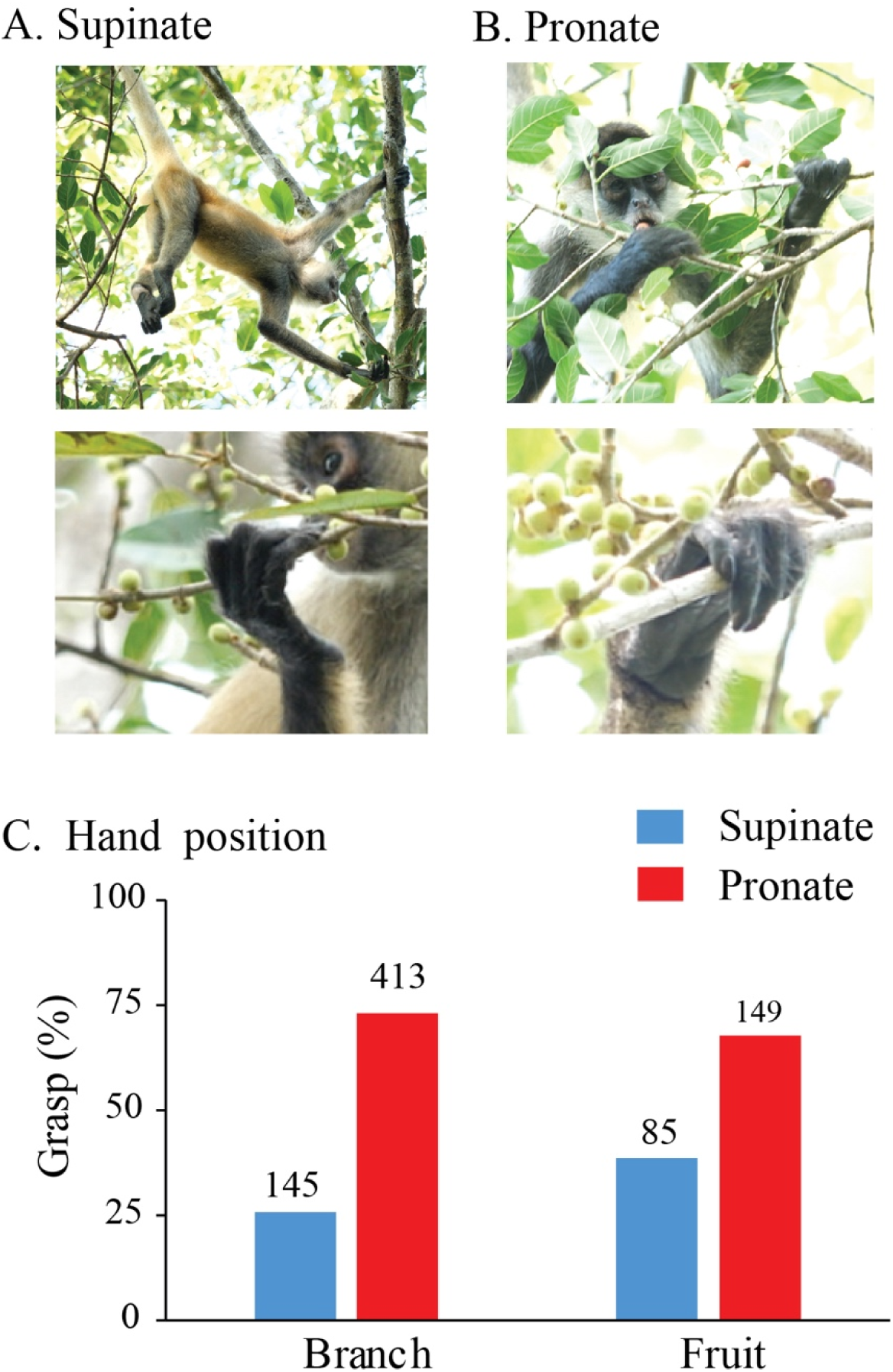
**A spider monkey called Kenya displays hand rotation associated with picking fruit or grasping branches containing fruit**. *Note.* A. The spider monkey reaches with a supinated hand to take a *Fictus cotinifolia* fruit item from a branch, as shown in whole body (top) and close-up (bottom) hand views. B. A pronated hand used when grasping a branch in a branch-withdraw to obtain the fruit *Fictus ovalis*, as shown in whole body (top) and close-up (bottom) views. C. The relative use of hand supination and pronation by the spider monkeys associated with branch-withdraw and grasp-withdraw movements showing that there is a preference for pronation-related grasp movements.

#### Hand grasps and mouth presentation

Figure 11 (Video 8) illustrates a representative hand posture used by a spider monkey grasping a fruit item and a representative hand posture the monkey used for presenting that fruit item to the mouth. The details of how a grasp was made were obtained from the observation of 66 grasps of small fruit items (<1.5 cm) and 45 large fruit items (>1.5 cm). All grasps were made with the radial side of the hand, with smaller fruit items held between the second digit (index) and the thenar side of the palm (Figure 11B) and larger food items additionally held with the third digit holding the food item against the palm. A grasped food item was always visible when the hand was viewed from its radial side (Figure 11B), as is also apparent in video of Nelson et al (2024) for a spider monkey grasping a small food item from a tabletop. We have previously designated grasps made between select digits pads and the palm as precision-power grips (Whishaw et al, 2024). When a food item was grasped using a pronated hand, the hand was counter-rotated to fit the food item into the hand. When a food item was grasped using a supinated hand, the hand was positioned so that the food item was in contact with the palm before the fingers were closed for purchase. Food presentations to the mouth were nearly always made with the radial side of the hand brought adjacent to the mouth where a fruit item was taken by the incisors (Figure 11C, n = 989/993 observations).

**Figure 11.**
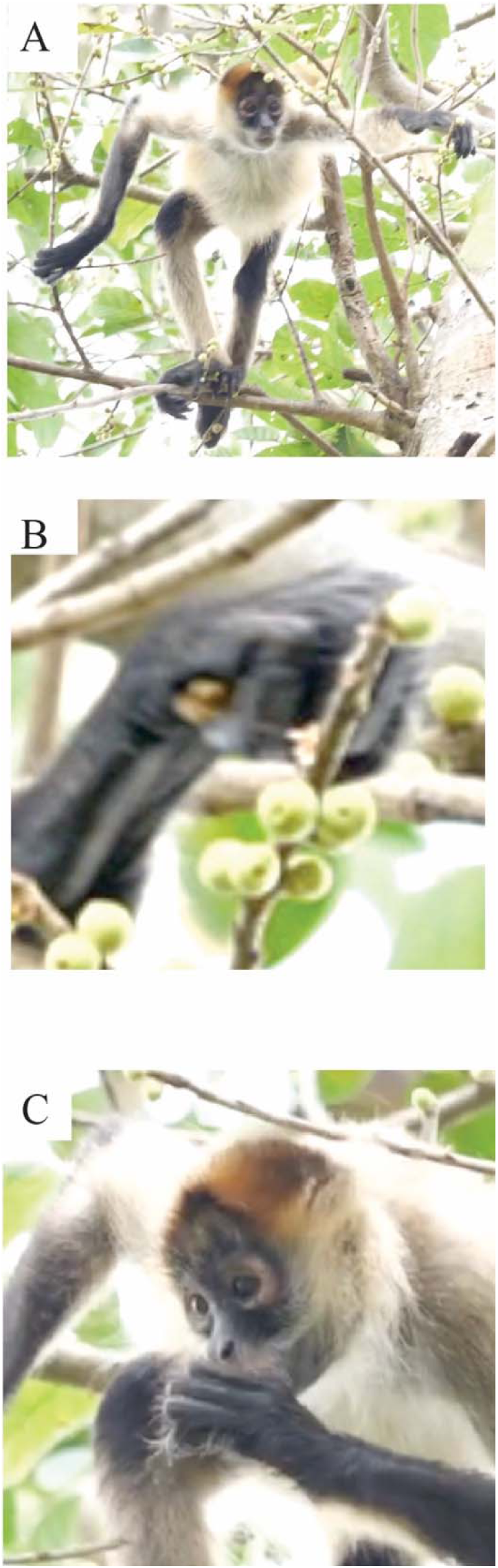
**Grasp-withdraw hand posture used for fruit picking and mouth placement the spider monkey Ucrania**. *Note*. A. Reaching for a fruit item of *Fictus cotinifolia*. B. Grasping fruit using a precision-power grip, in which the second digit holds the fruit against the radial side of the palm, with the fruit visible on the radial side of the palm. C. Placing fruit in the mouth with the radial side of the hand juxtaposed to the mouth and the hand in a vertically oriented posture.

Many rotational movements of the hand were associated with each transfer of a food item from the hand to the mouth (Figure 12). Figure 12A shows a large fruit item, *Simarouba glauca*, grasped by a spider monkey with opposition of the palm to digits 2 and 3, and Figure 12B shows the fruit presented to mouth by the radial side of the hand. Although a radial view of the hand showed that the fruit was visible, and so likely accessible for transfer from the radial side of the hand to the mouth, nevertheless, to assist in removing the fruit from the hand, the spider monkeys made rotational movements at the wrist (Figure 11C,D) and the upper arm (Figure 12 E,F) and also rotated the mouth to assist extraction of the food from the hand with the incisor teeth. These rotation movements were not counted but observed to occur in various configurations in all of 989 observations of food transfer to the incisors. Presentation of a food item to the side of the mouth so that it could be taken with the premolar teeth was observed on only three in hand-withdraws in one spider monkey eating the fruit *Guauma ulmifolia*.

**Figure 12.**
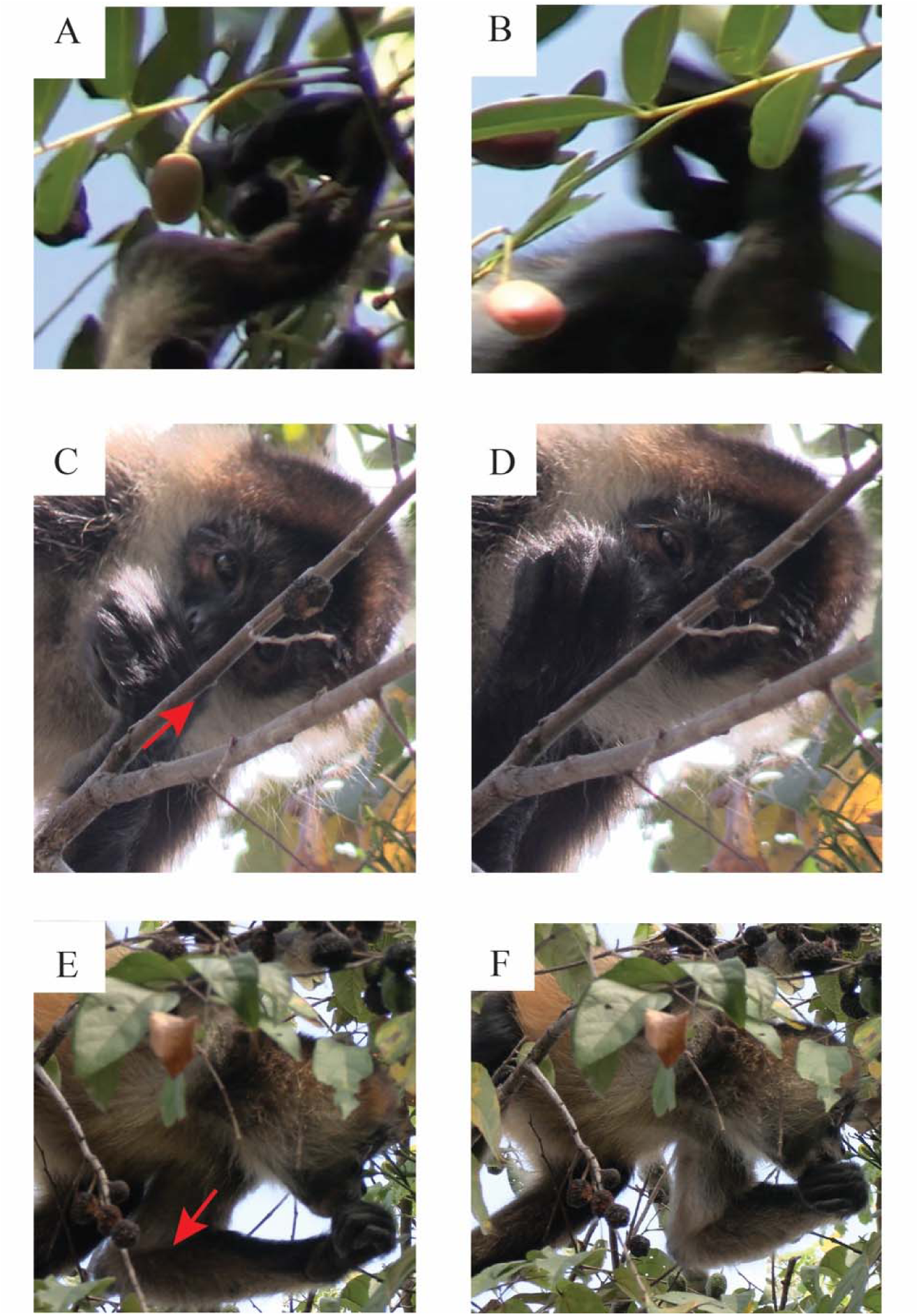
**Grasp-withdraw hand posture used for fruit picking and mouth placement by the spider monkey Bergen**. *Note*. A. Reaching for a fruit item of *Simarouba glauca*. B. The monkey uses a precision-power grip, in which the second digit holds the fruit against the radial side of the palm, with the fruit visible on the radial side of the palm. C. The monkey places the fruit in the mouth with the radial side of the hand juxtaposed to the mouth and the hand in a vertically oriented posture. C-D. To aid in extracting the food item from the hand, the spider monkey makes supination-pronation hand movements around the wrist, as shown by the arrow, that expose the fruit item to the mouth for grasping. E-F. To aid in extracting the food item from the hand, the spider monkey makes movements around the upper arm, as shown by the arrow, to change the vertical orientation of the hand to expose the fruit item to the mouth for grasping.

## Discussion

This study describes how individual spider monkeys in population of Costa Rica spider monkeys (*Ateles geoffroyi*) pick fruit while foraging in the upper tree canopy. The results provide insights into how the spider monkeys positional behavior, vision, and hand use support a reach-centered strategy of fruit harvesting. The spider monkey physique and reaching behavior suggest that they have evolved this reach phenotype to allow preferential access to even the most difficult-to-reach fruit. Phenotypic specialization in the reach in spider monkeys supports the idea that the two components of a reaching movement, the reach and the grasp, are distinct behavioral components of forelimb use in feeding that can differentially contribute to phenotypic behavioral variation. Our behavioral observations extend previous descriptions of the curious morphology of spider monkeys by describing how their form contributes to their phenotypic skills in fruit harvesting in contrast to the grasp phenotype of the sympatric capuchin monkey tree-canopy competitor.

Jeannerod’s (1981) dual visuomotor proposes that the reach of the forelimb is directed to the intrinsic (location) features of a target whereas the grasp shapes the hand in relation to the target’s intrinsic (size and shape) features. Whereas the theory has been extensively debated in experiments related to human forelimb use, it is relevant to questions related to the evolution of the reach and the grasp in primates (Karl and Whishaw, 2013; MacNeilage, 1987; Whishaw and Karl, 2019). Here we highlight features of the fruit-picking behavior of spider monkeys that suggest that they are specialized in the use of the reach. First, spider monkeys preferentially picked fruit by reaching for and hooking branches and pulling them in toward the mouth. Instead of grasping the fruit with their other hand, however, they then grasp the fruit from the branch with their mouth. That is, they use their reach to manipulate the branch as a tool or proto-tool to reel in fruit (St Amant and Horton, 2008; Seed et al., 2015). The branch-withdraw strategy was used for all fruit species but was most frequently used when harvesting smaller fruit items (see Figure 6). The branch-reach strategy was facilitated by their flexible rotatory movements around forelimb joints, which allowed them to pronate and supinate their hand to grasp and manipulate branches into a position optimal for the mouth grasping. Rosenberger et al. (2008) described spider monkey forelimb skeletal modifications that enable such flexibility in forelimb movement.

Second, tail prehension also supports the idea that spider monkeys are reach specialists. It has been observed that spider monkeys use tail support when feeding (Rosenberger et al., 2008; van Roosmalen, 1985; Youlatos et al., 2008), and the present quantification shows that up to half of their feeding featured tail support, with no differences related to sex or age. Even reaches in which the long axis of the body was in a horizontal position or in a sitting position could be associated with tail support as the spider monkeys leaned out to purchase a fruit item. Our definition of tail support was conservative, based on whether an animal might fall if the tail let go. Despite our conservative scoring of tail support, the spider monkeys were holding on to a branch with their tail on almost every reach, so it is possible that tail prehension was additionally adopted to assist with pending reaches and/or to provide positional comfort. The use of tail support appears to indicate that tail prehension of itself additionally contributes to the reach phenotype of the monkeys by allowing them to increase the extent of their reaches to grasp branches.

Third, the specialized morphology supporting mouth use is another line of evidence for spider monkeys as reach specialists. The spider monkeys use their mouth in much the same way that they might use a hand when reaching and grasping, in that they direct their mouth to a fruit target using vision and then grasp the fruit, assisted with touch cues, with their incisors processing. The spider monkey’s do not then sit up to eat but process the fruit while maintain the same posture. They transfer the fruit from the incisors to the molars, which chew the endocarp from fruit allowing it to be swallowed. With respect to fruit processing, inspection of the seeds in scat has shown that they are undamaged, which has suggesting to some investigators that spider monkeys do not chew (van Roosmalen,1985). Here we show that *Ateles geoffroyi*s always chews fruit, spits out the exocarp of some fruit such as for *Guazuma ulmifolia* and *Ficus cotinifolia*, and chews *Doliocarpus dentatus* leaving the exocarp attached to the stem. The ability of spider monkeys to reach, grasp and process fruit in rapid sequence while often suspended by their tail is additionally assisted by their flexibility of their upper body including their neck as has been described by Rosenberger et al. (2008). In all the use of the mouth in grasping and the rapid processing of fruit contributes to the efficiency of their reach-centered behavior.

Fourth, although spider monkeys do use their hands to pick fruit they appear to have difficulties in forming a grasp and in transferring fruit from the hand to the mouth. The SSR population of spider monkeys has no external thumb (Melin et al., 2022), and it is possible that this anatomy aids in hooking branches but impairs hand grasps and food transfer. The monkeys located the fruit visually but then visually disengaged and used touch cues to position the hand to grasp, a strategy that increased the time required to pick a fruit item, especially if fruit was small (Figure 4). Grasps were made between the pads of the second and third fingers and the palm with a precision-power grasp (Whishaw et al., 2024b), with the result that the grasped fruit was enveloped, but visible, in the radial side of the hand. Although the monkeys always brought the radial side of the hand to the mouth for food transfer, they then had to root the fruit out of the hand using hand and mouth rotational movements. The spider monkey’s difficulties in grasping seemed to be first related to positioning the fruit against the palm so that it could be pinned there by the fingers and then removed from the palm by the mouth, especially since simply opening the fingers presents a risk of dropping the fruit item. The difficulties in grasping and transferring fruit to the mouth might be secondary to their preferential use of their mouth to grasp fruit.

It has been proposed that the preferential use of the right hand by humans has an evolutionary history in which the left hand was specialized for the reach and the right hand was specialized for the grasp (MacNeilage, 1987). As a reach specialist, the spider monkey reach and grasp during natural foraging provides a good test of this idea. Nevertheless, we found that for natural foraging, spider monkey hand use for grasping branches or food did not feature a population asymmetry, although many animals displayed individual asymmetry as has been previously reported (Hook-Costigan et al., 1996; Motes Rodrigo et al., 2018; Nelson et al., 2015a; 2015b). In addition to reaching for branches or fruit, the spider monkeys could also hold a food item in hand and bring it repeatedly to the mouth for biting with the incisors or premolars, as do other simians including humans (Hirsche et al., 2022; Whishaw et al., 2025). This behavior was only observed when the spider monkeys ate *Guazuma ulmifolia*, a difficult-to-eat fruit item. Nevertheless, this behavior also did not feature a population handedness bias, as is found for an equivalent small human sample eating fruit in a similar way (Whishaw et al., 2025). Thus, the reach specialization of spider monkeys features ambidexterity and not the lateralization of that characterizes human food eating.

A reach-centered phenotype may provide spider monkeys with a foraging advantage over other fruit-eating primates, such as their sympatric tree-canopy competitor, the capuchin monkey, a species that can be described as a grasp-specialist (Christel & Fragaszy, 2000; Costello & Fragaszy, 1988; Melin et al., 2022; Spinozzi et al., 2004; Truppa et al., 2019, 2021; Whishaw et al., 2024ab). The contrast between spider monkey reach-centered and capuchin grasp-centered foraging may also represent an example of niche partitioning (Finke and Snyder, 2008; Schreier et al., 2009; Ungar, 1996). Although Chapman (1987) argued against niche partitioning based on the similar diets of *Ateles* and *Cebus,* we are suggesting that it is not diet that influences niche partitioning, but the differential behavior used for obtaining food. Future investigation could include direct comparisons of how the species-distinctive reaching strategies of spider monkeys and capuchin monkeys partition the canopy in relation to fruit acquisition. Future investigation might also consider whether the different foraging strategies of these primates influences the seeding of new trees.

In conclusion, it is relevant to consider whether a reach phenotype of spider monkeys is antecedent to some of their other behavior, including locomotion. Identifying the relative importance and order of selective agents that shaped morphological adaptations is an important but elusive goal. Suarez (2006; see also van Roosmalen, 1985) reports that the travel distance of spider monkeys is amongst the greatest in primates, which Youlatos et al. (2008) describe as featuring a forelimb reach strategy and prehensile tail assistance. The idea has been raised that tail-assisted suspensory reaching was an evolutionary precursor to tail-assisted locomotion (Rosenberger et al., 2008; Youlatos et al, 2008). Future research on other atelid primates (e.g., genera *Alouatta*, *Brachyteles*, and *Lagothrix*) with differing diets, and on taxa with convergent morphological adaptations, such as gibbons, may shed light on the influence of reach vs. grasp phenotypes on this fascinating area of research. Finally, it is reported that spider monkeys have a large brain, as do other fruit eating primates. The contribution of spider monkey morphology and reach-centered feeding could be further examined with anatomical analyse, including analyses of brain neocortical and subcortical motor pathways as has been done for capuchin monkeys (Bortoff and Strick, 1993; Strick et al., 2021).

## Acknowledgements

We thank R. Blanco Segura and M.M. Chavarria and other staff of the Área de Conservación Guanacaste and Ministerio de Ambiente y Energía. Behaviorral data were collected under research permits

## Author Contributions

IQW: Conceptualization, Methodology, Formal analysis, Investigation, Resources, Data curation, Writing-Original Draft, Writing-Review & Editing, Visualization, Supervision, Project administration; JD: Formal analysis; PRG Investigation, Writing-Review & Editing; EMC: Investigation, Writing-Review & Editing; MAM: Investigation, Writing-Review & Editing; FAC Writing-Review & Editing, Data curation, Funding acquisition; FA: Writing-Review & Editing, Data curation, Funding acquisition; ADM: Methodology, Data curation, Writing-Review & Editing, Supervision, Project administration, Funding acquisition

## Video captions

**Video 1.** Positional behavior facilitated by tail grasping. A spider monkey called Estados Unidos moves from a head down vertical posture to an almost horizontal position while using both hands to gather fruit-bearing branches to bring fruit to the mouth for mouth-associated fruit picking. Note: Hind limbs are providing minimal support. (30f/sec).

**Video 2**. Fruit picking and sucking. A spider monkey called Kenya takes the fruit *Doliocarpus dentatus* by mouth. The first mouth grasp involves picking the fruit, chewing, and spitting out the shell. The second and third mouth grasps involve taking the fruit in the mouth and chewing/sucking out the pulp while leaving the shell attached to the branch. (Slow-motion, 10% of 30 f/sec speed).

**Video 3**. A spider monkey named UN makes a grasp-withdraw of a *Ficus cotinifolia* fruit item. Note: A. Visual disengage occurs well before the contact of the hand with the fruit item. B. Hand adjustments associated with tactile mediated grasp of the fruit item. C. Presentation of the fruit to the mouth with the radial side of the hand. (Slow-motion, 10% of 30 f/sec speed).

**Video 4**. A spider monkey named UN makes a branch-withdraw of a *Ficus cotinifolia* fruit item. Note: A. Blink-associated disengage during the reach for the food item. B. The lips and incisors both participate in picking the fruit. C. Mouth adjustments associated with tactile-mediated grasp of the fruit item. (Slow-motion, 10% of 30 f/sec speed).

**Video 5**. A spider monkey called Ajka displays an inhand-withdraw in which the fruit *Guazuma ulmifolia* is held in hand and then brought to the mouth for chewing to remove the shell. Note: A. The inhand-withdraw movement consisted of movement components: A. Directing gaze to the fruit item and withdrawing the hand toward the mouth; B. Disengaging gaze with a blink when completing the withdrawal of the hand toward the mouth; and C. Grasping the fruit with the mouth from the radial side of the hand and releasing the fruit to the mouth for biting with the incisors. (Slow-motion, 10% of 30 f/sec speed).

**Video 6**. A spider monkey called Kenya displays fruit chewing during successive grasp-withdraw and branch-withdraw movements for the fruit *Ficus ovalis*. (Slow-motion, 10% of 30 f/sec speed).

**Video 7**. A spider-monkey called Poltava reaches for the fruit *Simarouba glauca* by mouth. After chewing the fruit, the spider-monkey raises its head to swallow the fruit (Slow-motion, 10% of 30 f/sec speed).

**Video 8**. A spider monkey called UN makes a grasp of the fruit item *Ficus cotinifolia*. Note A. Touch is used to place the radial side of the palm (thenar pads) on the fruit. B. The hand is counter-rotated to bring the second digit into position to grasp the fig against the palm. C. The fig is visible on the radial side of the hand. (Slow-motion, 10% of 30 f/sec speed).

